# The Effects of Ketamine on Methamphetamine Withdrawal-Induced Anxiety and Drug-Seeking Behaviors in the Rat

**DOI:** 10.1101/2025.06.24.661345

**Authors:** Marco German Ghilotti, Ricardo Petrilli Fortuna, Kofi Osei-Abrefa Ayensu, Danielle Stern, Ellen M. Unterwald

## Abstract

**Background:** The use of methamphetamine has continued to rise in the US. In addition to facilitating dopamine neurotransmission, methamphetamine indirectly increases glutamate release which activates N-methyl-D-aspartate receptors (NMDARs). Ketamine is a noncompetitive NMDAR antagonist. Ketamine also has actions on α-amino-3-hydroxy-5-methyl-4-isoxazolepropionic acid receptors (AMPARs) and promotes synaptogenesis. Thus, we hypothesized that ketamine may be a potential therapeutic to reduce methamphetamine-seeking behaviors and associated negative affect in a rat model.

**Methods:** Male and female rats underwent methamphetamine or saline intravenous self-administration for 10 sessions, followed by extinction training. Rats received ketamine or saline treatment either prior to 10 daily extinction sessions or only prior to the last extinction session. Anxiety-like behaviors were measured 24 hours after extinction, followed by cue-induced and drug-primed reinstatement two and six days later respectively.

**Results:** Methamphetamine withdrawal increased anxiety-like behaviors in male rats on the elevated plus maze test compared to rats that self-administered saline. Moreover, anxiety-like behaviors were significantly attenuated by daily ketamine treatment during extinction. Drug-primed but not cue-induced reinstatement, tested six days after the last extinction session, was significantly attenuated in male rats that received ten or one ketamine treatments during extinction compared with rats receiving vehicle during extinction. Ketamine was ineffective in female rats in reducing cue-induced or drug-primed reinstatement.

**Conclusions:** Ketamine may confer sex-specific benefits during methamphetamine withdrawal and relapse vulnerability, particularly by reducing anxiety-like behaviors and attenuating drug-primed reinstatement in males. These results support the potential of ketamine as a targeted adjunct therapy during early methamphetamine abstinence in males.

## 1. Introduction

Clinical manifestations of methamphetamine use include anxiety, cognitive dysfunction, and exacerbation of pre-existing psychiatric symptoms (McKetin et al., 2016; Paulus and Stewart, 2020). In 2021, approximately 2.5 million people aged 12 and older reported using methamphetamine, with around 1.6 million developing methamphetamine use disorder (MUD) (*Key Substance Use and Mental Health Indicators in the United States: Results from the 2021 National Survey on Drug Use and Health*, 2021). No pharmacological treatment has proven effective in treating MUD (“NIDA. 2021, April 13. What treatments are effective for people who misuse methamphetamine?,” n.d.).

The pharmacological action of methamphetamine is characterized by a prolonged rise in synaptic monoamine levels caused by the modulation of pre-synaptic neurotransmission. Methamphetamine-stimulated dopamine release is crucial for its reinforcing and neurobehavioral effects (Kogan et al., 1976; Parsegian and See, 2014;Fleckenstein et al., 2007). Acute or repeated exposure to methamphetamine also dysregulates glutamate transmission. Methamphetamine induces glutamate release and activation of NMDA receptors, while methamphetamine abstinence is associated with reduced basal glutamate in both rodent models and persons with MUD (Shrestha et al., 2022; Mark et al., 2004;Parsegian and See, 2014; Pena-Bravo et al., 2019; Kalivas and Volkow, 2005;Hámor et al., 2023; Nordahl et al., 2003). Reduced glutamate during abstinence is a result of increased excitotoxic glutamate activity during active methamphetamine exposure and the associated neuronal injury (Shrestha et al., 2022; Szumlinski et al., 2017;Smith et al., 2008). A prominent sequel to chronic methamphetamine is the sudden surge of glutamate that occurs after a period of abstinence in response to re-exposure to methamphetamine or methamphetamine-associated cues (Mark et al., 2004; Parsegian and See, 2014). Maintaining a balance of glutamate is crucial for the processes of neuroplasticity and synaptogenesis (Mattson, 2008). As such, disrupted glutamate during methamphetamine use likely contributes to dysregulated neuroplasticity and associated neurocognitive and behavioral deficits.

Given the impact of methamphetamine on the glutamate system, this study investigated the potential of ketamine to mitigate the adverse behavioral effects of chronic methamphetamine exposure in a rat model of methamphetamine self-administration. Ketamine is a non-competitive antagonist of NMDA receptors. It can act on three distinct NMDA receptor populations: receptors on GABA interneurons, on dopaminergic pre-synaptic neurons, and on neurons post-synaptic to dopamine terminals (Zanos and Gould, 2018). The effects of ketamine are numerous and include disinhibiting glutamate transmission, blocking NMDA receptor currents in GABAergic interneurons, raising extracellular glutamate concentrations in the prefrontal cortex (PFC), and activating AMPA receptors (Moghaddam et al., 1997; Zarate and Manji, 2008; Aleksandrova et al., 2017; Suzuki et al., 2023; Zaytseva et al., 2023). Ketamine administration and antidepressant-like effects are associated with a higher AMPA/NMDA receptor density ratio in rat hippocampus (Tizabi et al., 2012). Enhanced glutamatergic activity mediated by AMPA receptors may be responsible for increased synaptic potentiation and activation of early neuroplastic genes observed after ketamine (Machado-Vieira et al., 2009). Because of the ability of ketamine to modulate glutamate transmission, the objective of this study was to test the effectiveness of ketamine in reducing methamphetamine-seeking and anxiety-like behaviors produced by methamphetamine withdrawal in a rat model.

## 2. Materials and Methods

### 2.1 Animals

Male and female Sprague-Dawley rats from Charles Rivers Laboratories (Wilmington, MA), 8 weeks old upon arrival, acclimated to the facility for one week. Rats were maintained on a reverse 12-hour light/dark cycle (lights off at 9:00 AM) and food and water were provided ad libitum. Testing occurred during the dark phase. All methods were approved by the Institutional Animal Care and Use Committee of Temple University and complied with the NIH Guide for the Care and Use of Laboratory Animals.

### 2.2 Drugs

Methamphetamine (generously obtained from the NIDA Drug Supply Program) and (+/-)-ketamine hydrochloride (Sigma Aldrich Co, St. Louis, MO #K2753) were dissolved in sterile saline.

### 2.3 Jugular Vein Catheter Implantation Surgery

Rats were anesthetized with isoflurane, and polyurethane catheters (SAI Infusions, Lake Villa, IL) were implanted into the jugular vein as previously described (Connelly et al., 2020). Catheters were threaded subcutaneously over the scapula and affixed to a port mounted in the back (Plastics One, Roanoke, VA). Post-operation care included meloxicam as an analgesic and flushing the catheter twice daily with 0.2ml of heparinized saline (100 USP/ml) and enrofloxacin (0.225 mg/kg)(Li et al., 2022). Rats were allowed 7-14 days of recovery before beginning self-administration.

### 2.4 Methamphetamine Self-Administration

<u>Acquisition</u>: Self-administration was carried out in standard operant chambers (Med Associates, St. Albans, VT) as described previously (Connelly et al., 2020). Each 2-hr session began with the activation of a house light and the extension of two levers. Presses on the active lever activated a syringe pump to deliver 0.05mg/kg/inf of methamphetamine or saline according to a Fixed-Ratio 1 (FR1) reinforcement schedule. Each rewarded lever press illuminated a cue light in conjugation with a tone. There was a 20-sec time-out period after each infusion. Lever presses on the inactive lever were logged but had no repercussions. After reaching an initial acquisition criteria of ≥10 infusions, rats completed ten sessions of methamphetamine self-administration; saline self-administration occurred during 10-12 sessions under identical conditions. <u>Extinction</u> training began 2 days later wherein rats were pretreated with ketamine (0, 5, or 10 mg/kg ip) and, 30 minutes later, placed into the chambers for 2-hrs. Two ketamine injection protocols were tested: (1) ketamine, 5 mg/kg ip, prior to the 10 daily extinction sessions, or (2) one ketamine, 10 mg/kg ip, injection prior to the last extinction session only. During extinction, lever presses had no scheduled consequences (i.e., no infusion or cues). Extinction lasted for 10 sessions. <u>Reinstatement</u>: Through cue- and drug-primed (+cue) reinstatement, return to drug-seeking behavior was assessed. Cued reinstatement was tested 48-hrs after the last extinction session, wherein active lever presses resulted in the delivery of the light and tone (no drug delivery). Rats returned to extinction for 3 sessions to ensure extinguished responding. For the drug-primed (+cue) reinstatement test, methamphetamine (1mg/kg ip) was injected immediately before the session. Active lever pressing was accompanied by a light and a tone with no drug delivery. Cue- and drug-primed reinstatement tests both lasted 2-hrs. Inactive and active lever presses were recorded and analyzed.

### 2.6 Anxiety Testing

Anxiety-like behavior was measured 24-hrs after the last extinction session using the elevated plus maze (EPM) as previously described (Denny et al., 2021; Perrine et al., 2008). Rats were placed in the center of the EPM, and behaviors were recorded for 10-minClick or tap here to enter text.. The number of entries and time spent on each arm of the EPM were scored from the videos by two investigators, at least one was blind to the experimental group. The percentage of time spent in open arms (time in open arms/total open + closed arm time x 100) and the percentage of open arm entries (open arm entries/total arm entries x 100) served as indicators of anxiety-like behaviors.

### 2.7 Data Analysis

GraphPad Prism v10.0 software was used for statistical analysis. Unpaired two-tailed t-tests were used to assess significant diHerences between the two groups on reinstatement. Two-way ANOVAs followed by Tukey’s post hoc tests were performed on self-administration and EPM data. Individual data points are shown along with means ± SEMs. P<0.05 was deemed significant.

## 3. Results

The experimental timeline is shown in Fig 1A, during which rats self-administered methamphetamine (0.05 mg/kg/infusion) or saline for 10 days, followed by 10 extinction sessions with or without ketamine treatment. Rats that self-administered methamphetamine had higher active lever presses than rats that self-administered saline (Fig 1B). Two methamphetamine groups had similar active lever presses (Fig 1B) and intake (Fig 1C) during acquisition. During extinction, there was no significant difference in active lever presses between rats that received ketamine prior to each extinction session compared to those that received saline (Fig 1B), indicating that ketamine did not alter the rate of extinction.

**Figure 1:**
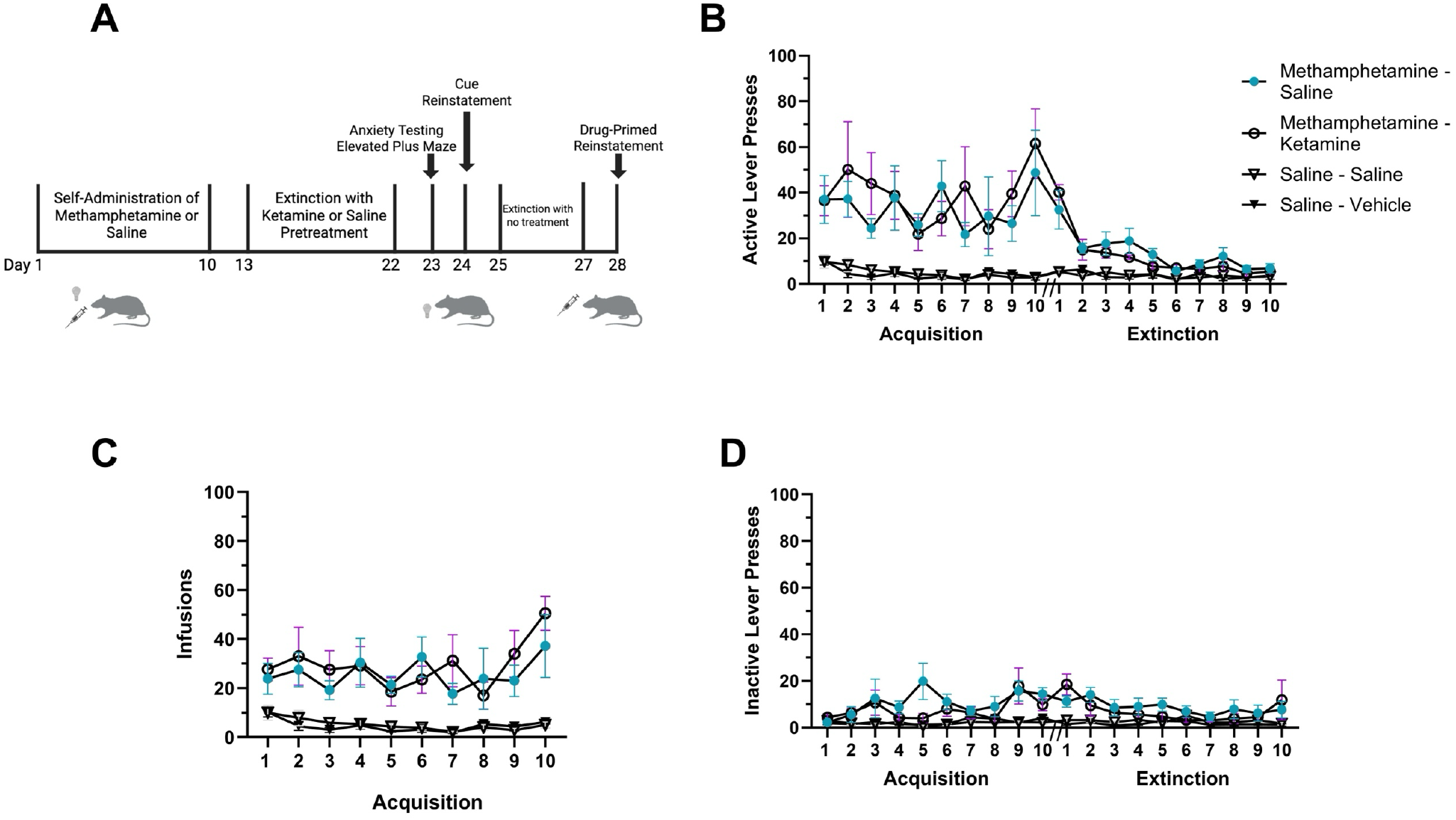
Methamphetamine self-administration and ketamine treatment during extinction. **1A:** Experimental Timeline, 10-day ketamine treatment in male rats. **1B:** Active lever presses for methamphetamine (0.05 mg/kg/inf) or saline are shown over 10 sessions of FR1 self-administration, followed by 10 extinction sessions. Ketamine (5 mg/kg ip) treatment occurred 30-min prior to each extinction session and did not affect responding during extinction. **1C**: Methamphetamine infusions earned are shown and did not differ between groups. **1D:** Inactive lever presses were similar across experimental groups. Data are expressed as means +/- SEMs; N=6-7.

Ketamine treatment during extinction training had no effect on cue-induced reinstatement tested 48-hrs after the last ketamine injection (t-test, t=1.602, df=11, p=0.1374; Fig 2A). However, drug-primed reinstatement was significantly attenuated in rats receiving ketamine during extinction as compared with saline-injected rats (t-test, t=3.468, df=11, p=0.0053; Fig 2B). Notably, drug-primed reinstatement occurred six days after the last ketamine injection.

**Figure 2:**
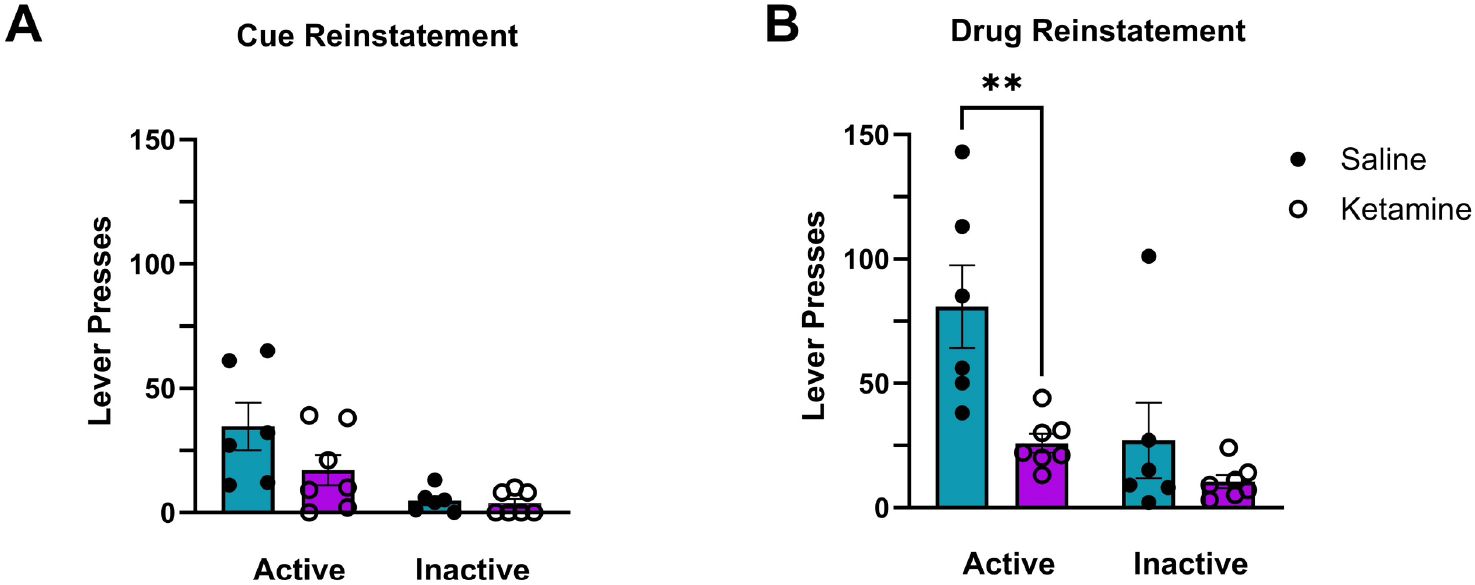
Ketamine treatment during extinction training reduced drug-primed reinstatement. **2A:** Responding on active and inactive levers during cue reinstatement for male rats treated with ketamine or saline daily during extinction training. **2B:** Ketamine reduced drug-primed reinstatement. Responding on active and inactive levers are shown for male rats treated with ketamine or saline during extinction training. (**p=0.0053; Means +/- SEM; N=6-7).

Anxiety-like behaviors were assessed 24-hrs after extinction training (two weeks after cessation of methamphetamine self-administration). Two-way ANOVA of percent open arm time on the EPM revealed a significant interaction (Fig 3A; F(1,20)=16.4, p=0.0006) between main effects of ketamine/saline (F(1,20)= 5.620, p=0.0279) and methamphetamine/saline (F(1,20)= 27.97, p<0.0001). Likewise, percent open arm entries (Fig 4B) showed significant interaction and main effects (Interaction: F(1,20)= 8.426, p=0.0088; ketamine: F(1,20)= 8.210, p=0.0096; methamphetamine: F(1,20)= 19.70, p=0.0003). Post-hoc analyses demonstrated that rats that self-administered methamphetamine and received vehicle injections during extinction exhibited higher anxiety-like behaviors with significantly lower open arm time (p<0.0001) and open arm entries (p<0.0001) compared to rats that self-administered saline. Methamphetamine withdrawal-induced anxiety was significantly attenuated by ketamine treatment during extinction. Rats self-administering methamphetamine and treated with ketamine had significantly less anxiety than those treated with vehicle (p<0.01), and open arm time and entries did not differ from controls, demonstrating that ketamine reduced anxiety-like behaviors produced by methamphetamine withdrawal (Fig 3). Ketamine had no effect on EPM performance in rats self-administering saline (Figs 3A and B).

**Figure 3:**
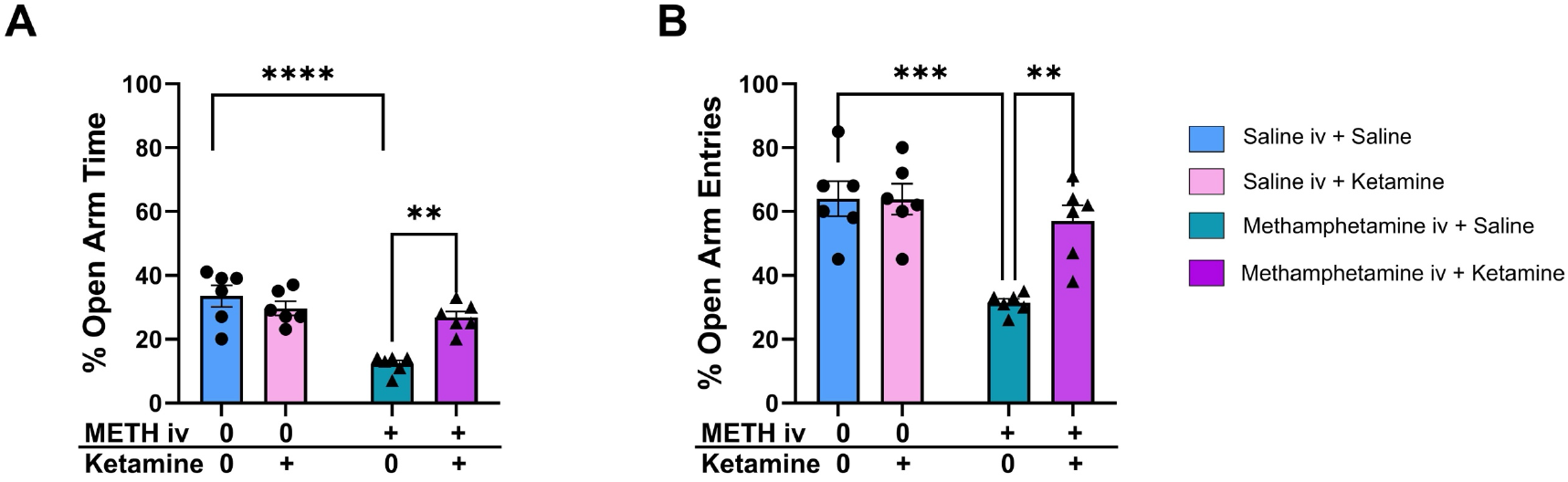
Ketamine attenuated methamphetamine withdrawal-induced anxiety. EPM testing occurred 24-hr after the last extinction session. Rats that self-administered methamphetamine had lower open arm time (**3A**) and entries (**3B**) compared with rats that self-administered saline. Ketamine administration during extinction training attenuated the reduction of open arm time and open arm entries produced by methamphetamine withdrawal, indicating an anxiolytic effect of ketamine. Two-way ANOVA with Tukey’s multiple comparisons test (****p=<0.0001, **p=0.0011, ***p=0.0002; N=6-7/group).

**Figure 4:**
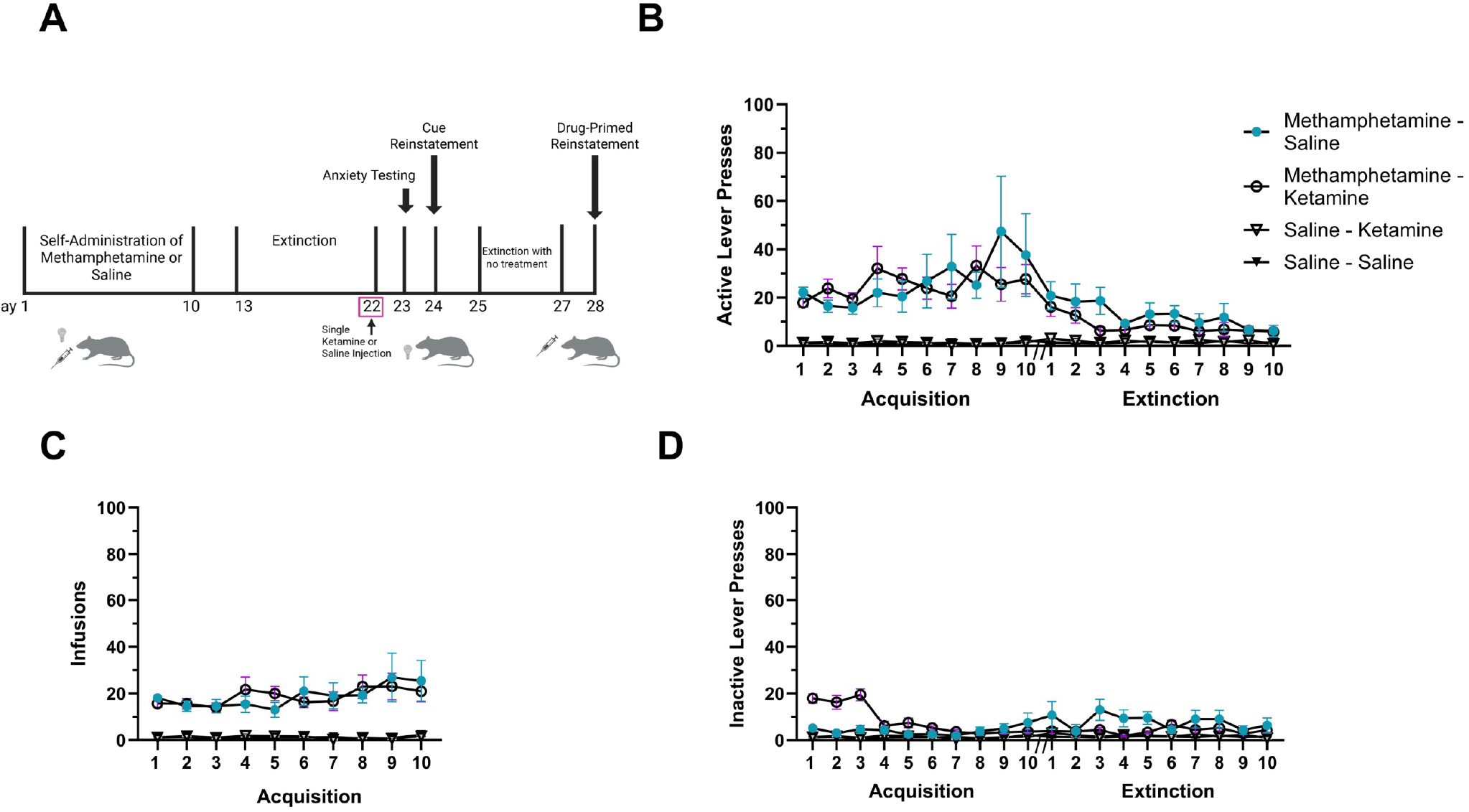
Methamphetamine self-administration and extinction. **4A:** Experimental Timeline 2: 1-day ketamine treatment with male rats. **4B:** Active lever presses for methamphetamine (0.05 mg/kg/infusion) or saline during 10 acquisition sessions, followed by 10 extinction sessions are shown. Ketamine (10 mg/kg) was administered prior to the last extinction session only. **4C:** The number of methamphetamine infusions earned was the same for the two experimental groups. **4D:** Inactive lever presses did not differ between treatment conditions. Data are expressed as means +/- SEMs; N=10-12 rats/group.

The next experiment tested whether a single ketamine injection would be sufficient to attenuate reinstatement. Male rats self-administered methamphetamine or saline as before and then went through 10 extinction sessions. Rats received ketamine (10 mg/kg) or saline prior to the last extinction session, as shown in the experimental timeline (Figure 4A). There were no significant differences in active lever presses (Fig 4B), inactive lever presses (Fig 4D), methamphetamine infusions (Fig 4C), or responding during extinction (Fig 4B) between groups, as expected.

Cue-induced reinstatement was tested 48-hr after extinction; no significant difference in active lever presses was found between rats receiving ketamine or saline on the last extinction day (Fig 5A; t-test, t=1.946, df=22, p=0.0646). However, rats treated with ketamine had significantly lower active lever presses than those treated with saline during drug-primed (+cue) reinstatement (Fig 5B; t-test, t= 2.971, df= 20,p=0.0076). Inactive lever responses were not different between groups (Fig 5A and 5B). Thus, both ketamine treatment regimens (eg, 1 or 10 days) suppressed drug-seeking behaviors in the drug-primed (+cue) reinstatement test but not in cue-alone reinstatement.

**Figure 5:**
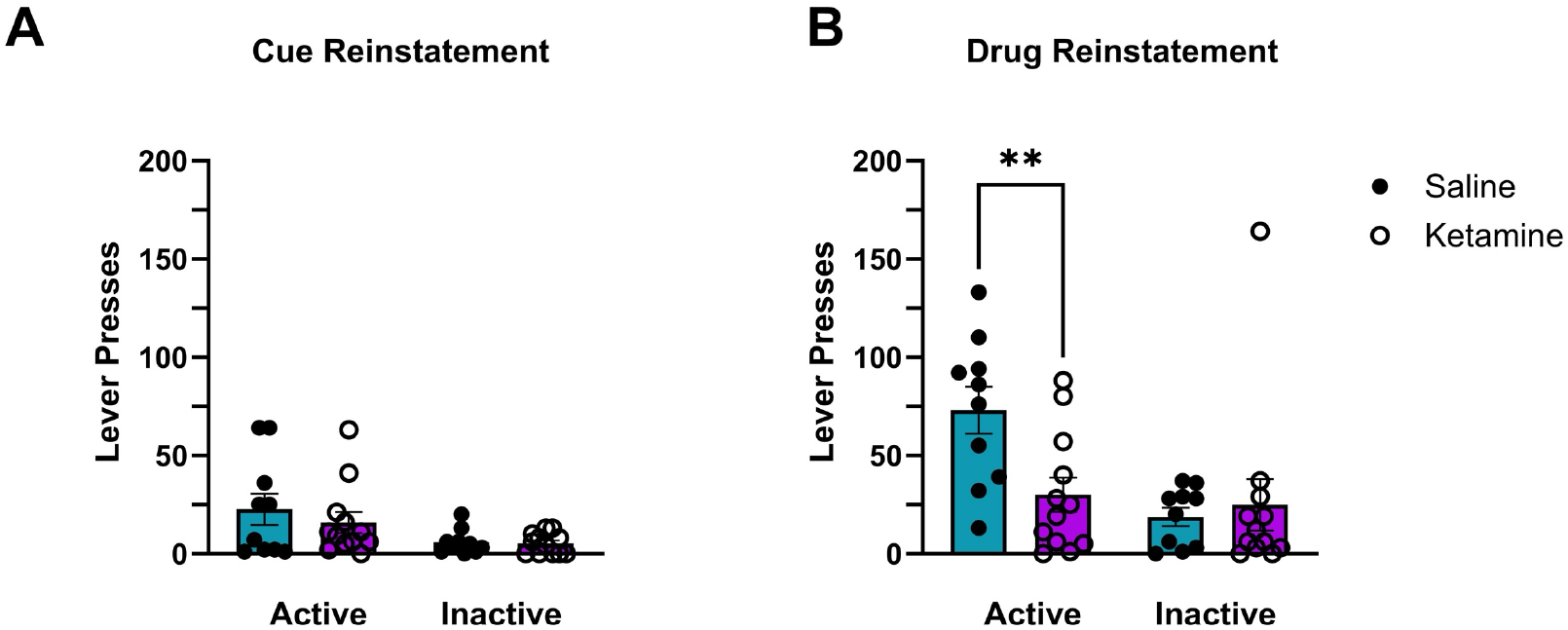
Ketamine treatment on extinction day 10 reduced drug-primed reinstatement. **5A**: Ketamine had no effect on active or inactive lever presses during cue-induced reinstatement. **5B:** Male fats treated with one injection of ketamine had lower active lever responses during drug-primed reinstatement compared with saline controls (**p=0.0076, N=10-12/group).

The next experiment followed the timeline outlined in Figure 6A using female rats. Female rats self-administered methamphetamine (0.05mg/kg/infusion) or saline for 10 days, followed by 10 extinction sessions (Fig 6B). During the extinction phase, female rats received a ketamine (5 mg/kg ip) or saline pretreatment 30 mins prior to each extinction session, totaling 10 ketamine or saline injections. Ketamine had no effect on active lever presses during the extinction phase (Fig 6B).

**Figure 6:**
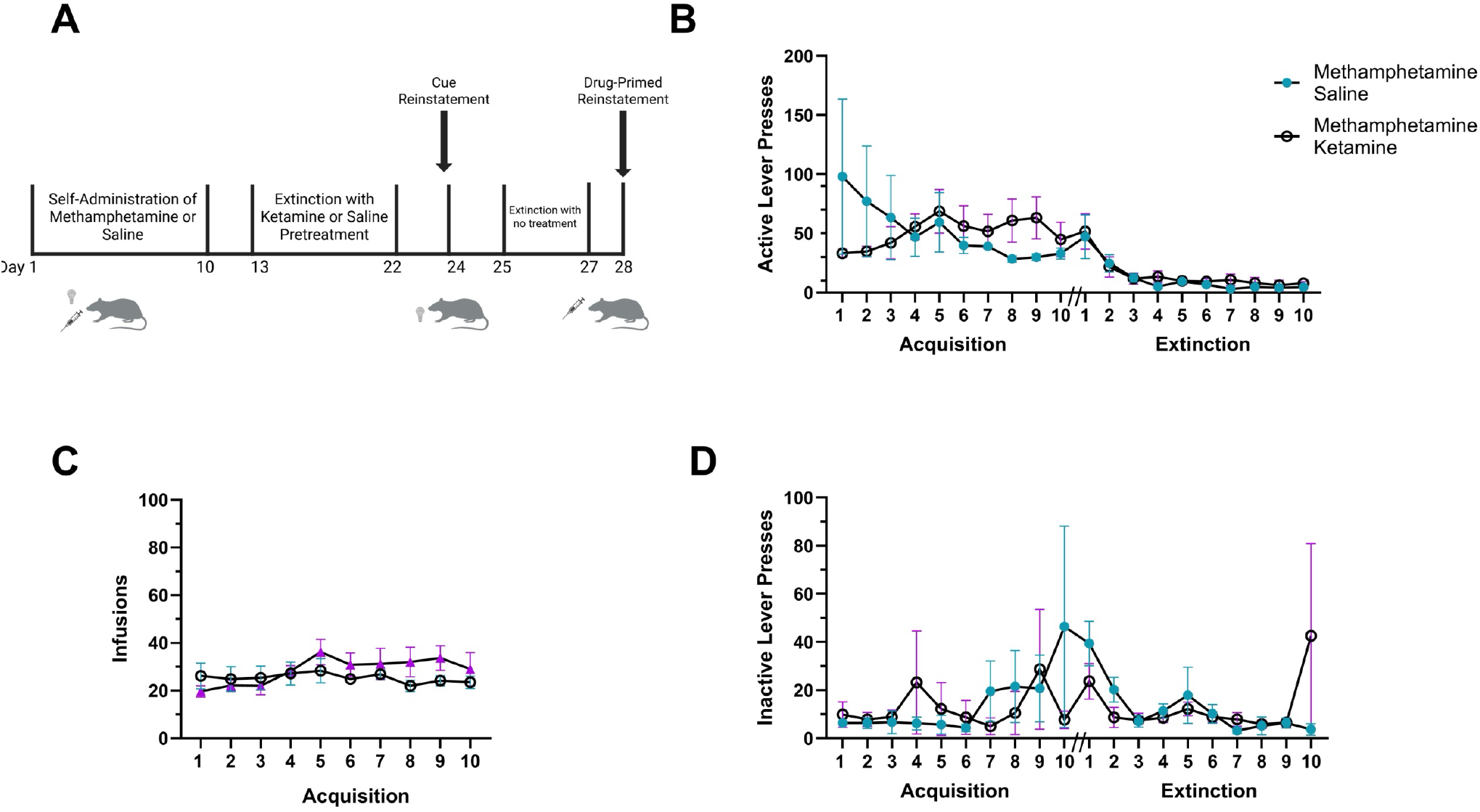
Methamphetamine self-administration in female rats. **6A:** Experimental Timeline 3: 10-day ketamine treatment. **6B:** Active lever presses for methamphetamine (0.05 mg/kg/inf) or saline are shown over 10 sessions of FR1 acquisition, followed by 10 extinction sessions. Ketamine (5 mg/kg ip) treatment occurred 30-min prior to each extinction session. There were no significant difference of active lever during either acquisition or extinction. **6C**: Methamphetamine infusions earned are shown and did not differ between groups. **6D:** Inactive lever presses were similar across treatment groups. Data are shown as means +/- SEMs; N=5-6/group.

Cue-induced reinstatement was tested 48-hrs after the last extinction session. Ketamine treatment during extinction has no significant effect on cue-induced reinstatement (t-test, t=0.5959, df=9, p=0.5659; Fig 7A). Likewise, drug-primed (+cue) reinstatement was not significantly different between ketamine and saline treatment groups (t-test, t=0.4775, df=8, p=0.6458; Fig 7B). Thus, daily ketamine treatment during extinction reduced methamphetamine-primed reinstatement in male rats but was ineffective in females.

**Figure 7:**
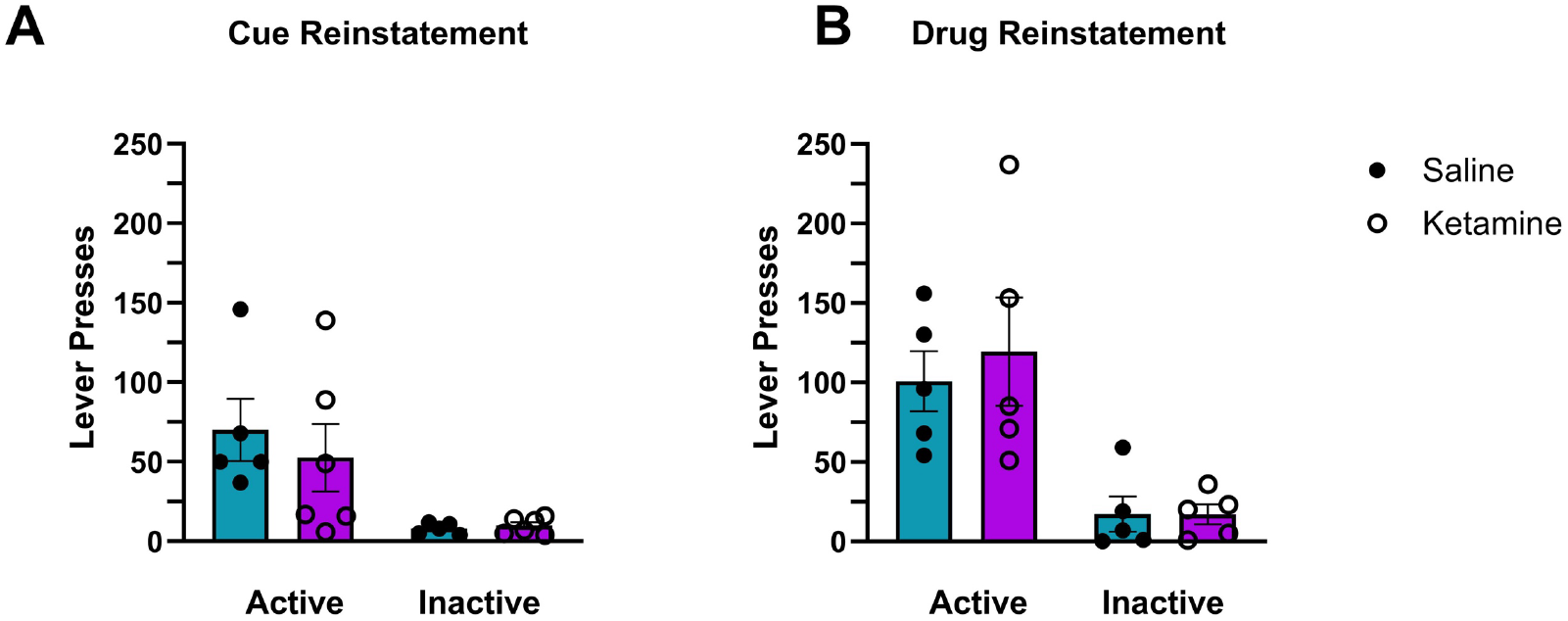
Ketamine treatment did not affect reinstatement in female rats. **7A**: Ketamine (5 mg/kg ip) administered prior to each extinction session had no effect on active or inactive lever pushes during cue-reinstatement (p=0.565). **7B:** Ketamine treatment had no impact on responding during drug-primed reinstatement in female rats (p=0.645). N= 5-6/group.

## 4. Discussion

The use of methamphetamine is a prominent public health problem (*Key Substance Use and Mental Health Indicators in the United States: Results from the 2021 National Survey on Drug Use and Health*, 2021). Repeated misuse of methamphetamine has the potential to lead to the development of a methamphetamine use disorder (MUD). Relapse rates are high, with an estimated 75% of people with MUD returning to drug use within 5 years of treatment (Paulus and Stewart, 2020). There are no FDA-approved medications for MUD, and as such, new treatment strategies are needed. This study aimed to provide critical preclinical evidence for the efficacy of ketamine in attenuating maladaptive behaviors produced by methamphetamine exposure.

Maintaining a balance of normal glutamate function is crucial for the processes of neuroplasticity and synaptogenesis. It is hypothesized that addiction, including to methamphetamine, is linked to persistent dysregulation of glutamate transmission, particularly in the prefrontal cortex (PFC) to the nucleus accumbens (NAc) circuit, leading to decreased synaptic plasticity and the development of maladaptive behaviors (Kalivas and Volkow, 2005; Kalivas et al., 2009). Acute or repeated methamphetamine elevates glutamate levels in several regions of the forebrain, including the PFC, NAc, and hippocampus, while withdrawal from chronic methamphetamine leads to reductions of basal glutamate levels in NAc and PFC (Nordahl et al., 2003; Hámor et al., 2023; Mark et al., 2004; Parsegian and See, 2014; Pena-Bravo et al., 2019; Kalivas et al., 2009). Maladaptive neuroadaptations produced by repeated methamphetamine and withdrawal are still being uncovered. *In vitro* studies have shown that NMDA receptor abundance or function undergoes adaptive modifications after withdrawal from long-term methamphetamine exposure (Smith et al., 2008). Augmentation of NMDA-EPSPs in striatal neurons of rats after methamphetamine exposure and withdrawal has been demonstrated and is associated with decreased Mg2+ blockage of the NMDA receptor and enhanced excitotoxicity and behavioral sensitization (Moriguchi et al., 2002).

Since methamphetamine impacts glutamate transmission, we hypothesized that ketamine, an NMDA receptor antagonist, could be effective in restoring normal glutamate function and attenuating methamphetamine-induced maladaptive behaviors. Ketamine enhances extracellular glutamate concentrations in the PFC by blocking NMDA receptor currents in GABAergic interneurons, disinhibiting glutamate transmission (Moghaddam et al., 1997; Zarate and Manji, 2008). Moreover, ketamine can augment AMPA receptor activity. An elevation in the AMPA/NMDA receptor density ratio has been noted in the hippocampus of rats following ketamine administration (Tizabi et al., 2012). Enhanced glutamatergic activity mediated by AMPA receptors, rather than NMDA receptors, may account for the increased synaptic potentiation and activation of early neuroplastic genes found following drug exposure (Machado-Vieira et al., 2009).

Similar to the way ketamine is used to treat PTSD, the conceptual underpinning for its use for MUD calls for its administration in conjunction with therapy in a medical environment (Duek et al., 2023). Under such conditions in persons with PTSD, a single ketamine infusion lessens fear reactions for more than 30 days (Duek et al., 2023). Benefits of ketamine have been seen in individuals with opioid use disorder (Anna et al., 2024; Krupitsky et al., 2007); abstinence rates at one year are enhanced by either one- or three-monthly ketamine treatments (Krupitsky et al., 2007). A single ketamine infusion significantly reduced cocaine use by 67% and cocaine craving at time points of >24 hours post-infusion, shown by a human laboratory study that assessed behavioral shifts in the salience of cocaine now versus money later (Dakwar et al., 2017; Dakwar et al., 2019). In some subjects, this effect persisted for at least two weeks. The authors propose that downstream pro-plasticity and modulatory mechanisms of ketamine’s therapeutic activities are involved, as effectiveness was shown at time points when ketamine and its metabolites were no longer present in the body (Dakwar et al., 2017; Dakwar et al., 2019). Ketamine, therefore, may help rewrite dysfunctional memories and compulsive behaviors in MUD by promoting neuroplasticity.

Existing literature highlights sex differences in the behavioral and neurobiological effects of methamphetamine exposure. Some studies report that female rats escalate methamphetamine intake more rapidly during extended access sessions and exhibit greater reinstatement of methamphetamine-seeking behavior compared to males (Daiwile et al., 2022; Reichel et al., 2012; Takashima et al., 2018; Cox et al., 2013; Roth and Carroll, 2004), whereas others show females take less methamphetamine than males under FR conditions (Daiwile et al., 2022; Funke et al., 2023; Lewandowski et al., 2023; Nagy et al., 2024; Ruda-Kucerova et al., 2015). The data presented here show similar responding for and intake of methamphetamine during the FR1 acquisition phase for males and females. Prior work demonstrates memory deficits in both sexes following extended methamphetamine access, yet females showed distinct glutamatergic adaptations (Daiwile et al., 2022; Takashima et al., 2018). Specifically, females exhibit reduced baseline spontaneous glutamate release but greater evoked glutamate release in the PFC after methamphetamine self-administration (Giacometti and Barker, 2020; Pena-Bravo et al., 2019). The heightened excitatory neurotransmission is accompanied by enhanced NMDA receptor currents and upregulation of GluN2B-lacking NMDA receptors in the PFC, changes not observed in males (Daiwile et al., 2021; Pena-Bravo et al., 2019). Additionally, reinstatement of methamphetamine-seeking behavior increased CaMKII expression in the dentate gyrus of males, whereas females showed a higher GluN2A/2B ratio, further supporting sex-specific glutamatergic signaling alterations (Takashima et al., 2018). Females also displayed lower basal excitatory postsynaptic currents (sEPSCs) in the PFC but showed increased evoked EPSC amplitude and NMDA receptor currents after methamphetamine exposure (Pena-Bravo et al., 2019). The GluN2B antagonist Ro256981 reduced NMDA currents in males but not in females, suggesting differences in NMDA receptor subunit composition and function between sexes (Pena-Bravo et al., 2019). Together, these findings indicate that the impact of methamphetamine exposure is different in males and females and underscore the importance of considering sex as a biological variable in the neurobiological mechanisms underlying methamphetamine addiction and relapse.

Interestingly, sex differences in ketamine metabolism and mechanisms of action have been noted. Female rats exhibit higher concentrations of ketamine and its active metabolite, norketamine, in both plasma and brain shortly after administration compared to males, a difference attributed to slower clearance rates and longer half-lives in females (Saland and Kabbaj, 2018). In a study of depression-like behaviors induced by chronic social isolation, ketamine effectively reversed these behaviors in both male and female rats, but its impact on neuronal morphology was sex-specific (Sarkar and Kabbaj, 2016). In males, ketamine restored spine density and synaptic protein levels in the medial PFC, but these structural changes were not observed in females, suggesting underlying sex-specific mechanisms of the effect of ketamine on neuroplasticity (Sarkar and Kabbaj, 2016). The absence of ketamine efficacy in reducing methamphetamine-primed or cue-induced reinstatement in female rats reported herein may reflect sex-specific differences in the pharmacokinetics and/or neuroplasticity effects of ketamine. Saland et al. (Saland and Kabbaj, 2018) shows that although female rats have higher concentrations of ketamine and norketamine, these elevated levels do not necessarily translate to equivalent neurobiological effects. In males, ketamine restores spine density and synaptic protein expression in the medial PFC, a region critically involved in extinction learning and regulation of drug-seeking. However, these structural changes are absent in females, suggesting that ketamine fails to engage, or sufficiently modulate the synaptic remodeling pathways necessary for its anti-relapse effects in females (Saland and Kabbaj, 2018). Future studies should examine how ketamine affects circuit-level activity and plasticity markers in females during extinction and reinstatement, and whether adjunctive therapies could enhance its efficacy.

Withdrawal from amphetamine and methamphetamine is associated with heightened anxiety in both humans and rodent models (Barr et al., 2010; McGregor et al., 2005; Su et al., 2017; Vuong et al., 2010; Zorick et al., 2010). As anxiety can contribute to relapse to drug use, alleviating anxiety could be therapeutically advantageous. Methamphetamine withdrawal reliably increased anxiety-like behaviors in male rats, consistent with previous research linking psychostimulant withdrawal to heightened negative affect in rodents (Barr et al., 2010; Perrine et al., 2008; Vuong et al., 2010; Ru et al., 2019). The finding that daily ketamine administration during extinction significantly reduced these anxiety-like behaviors when tested 24 hours later supports the anxiolytic potential of ketamine, possibly via its known effects on glutamatergic transmission and neural plasticity in brain regions such as the prefrontal cortex and amygdala.

Interestingly, while ketamine treatment during extinction attenuated drug-primed reinstatement in male rats, it had no effect on cue-induced reinstatement in either sex. This dissociation suggests that ketamine may be more effective in targeting reinstatement driven by pharmacological triggers (i.e., drug-induced) rather than conditioned cues. This specificity has implications for treatment strategies targeting relapse, indicating that ketamine may help reduce vulnerability to drug-provoked relapse rather than cue-driven craving, at least in males. In contrast to males, female rats failed to show significant reductions in drug-seeking behaviors following ketamine treatment, regardless of the reinstatement modality (cue- or drug-induced). This sex-specific lack of response may be due to a variety of factors, including differences in neurobiological adaptations to either methamphetamine and ketamine, hormonal influences, or divergent engagement of neural circuits underlying extinction and reinstatement. For example, estrogen and progesterone modulate stress and reward pathways (Flores et al., 2020; Giacometti et al., 2022), potentially blunting or altering the efficacy of ketamine in females. A limitation to the current study is that only a single ketamine dose and administration schedule was tested in the female rats; it is possible that other dosing regimens could be more efficacious.

## 5. Conclusions

The data presented provides critical preclinical data to advance the study of the potential therapeutic effects of ketamine for MUD. It sets the stage for future mechanistic studies that examine the action of ketamine on glutamate dysregulation and synaptogenesis in the setting of methamphetamine-induced neuroadaptations and drug-seeking behaviors. The results also challenge the assumption of uniform efficacy of ketamine across sexes and point to the necessity of tailored approaches in both research and clinical contexts. Future investigations should explore sex-specific molecular targets and consider hormonal cycles in females, which could modulate both behavioral and structural responses to treatment.

## Acknowledgements

This work was supported in part by NIH/NIDA P30DA013429 (EMU). We would like to thank Mary McCafferty, Daniel Lopez, and Leon Passarelli-Roberts for experimental support.

## References

Aleksandrova, L.R., Phillips, A.G., Wang, Y.T., 2017. Antidepressant effects of ketamine and the roles of AMPA glutamate receptors and other mechanisms beyond NMDA receptor antagonism. Journal of Psychiatry and Neuroscience 42, 222–229. 10.1503/jpn.160175

Anna, O., Michael, A., Apostolakis, M., Mammadov, E., Mitka, A., Kalatta, M.A., Koumas, M., Georgiou, A., Chatzittofis, A., Panayiotou, G., Gergiou, P., Zarate, C.A., Zanos, P., 2024. Ketamine and hydroxynorketamine as novel pharmacotherapies for the treatment of Opioid-Use Disorders. Biol Psychiatry. 10.1016/j.biopsych.2024.09.008

Barr, J.L., Renner, K.J., Forster, G.L., 2010. Withdrawal from chronic amphetamine produces persistent anxiety-like behavior but temporally-limited reductions in monoamines and neurogenesis in the adult rat dentate gyrus. Neuropharmacology 59, 395–405. 10.1016/j.neuropharm.2010.05.011

Connelly, K.L., Wolsh, C.C., Barr, J.L., Bauder, M., Hausch, F., Unterwald, E.M., 2020. Sex differences in the effect of the FKBP5 inhibitor SAFit2 on anxiety and stress-induced reinstatement following cocaine self-administration. Neurobiol Stress 13, 100232. 10.1016/j.ynstr.2020.100232

Cox, B.M., Young, A.B., See, R.E., Reichel, C.M., 2013. Sex differences in methamphetamine seeking in rats: Impact of oxytocin. Psychoneuroendocrinology 38, 2343–2353. 10.1016/j.psyneuen.2013.05.005

Daiwile, A.P., Jayanthi, S., Cadet, J.L., 2021. Sex-and Brain Region-specific Changes in Gene Expression in Male and Female Rats as Consequences of Methamphetamine Self-administration and Abstinence. Neuroscience 452, 265–279. 10.1016/j.neuroscience.2020.11.025

Daiwile, A.P., Sullivan, P., Jayanthi, S., Goldstein, D.S., Cadet, J.L., 2022. Sex-Specific Alterations in Dopamine Metabolism in the Brain after Methamphetamine Self-Administration. Int J Mol Sci 23, 4353. 10.3390/ijms23084353

Dakwar, E., Hart, C.L., Levin, F.R., Nunes, E. V, Foltin, R.W., 2017. Cocaine self-administration disrupted by the N-methyl-D-aspartate receptor antagonist ketamine: a randomized, crossover trial. Mol Psychiatry 22, 76–81. 10.1038/mp.2016.39

Dakwar, E., Nunes, E. V., Hart, C.L., Foltin, R.W., Mathew, S.J., Carpenter, K.M., Choi, C.J. “Jean,” Basaraba, C.N., Pavlicova, M., Levin, F.R., 2019. A Single Ketamine Infusion Combined With Mindfulness-Based Behavioral Modification to Treat Cocaine Dependence: A Randomized Clinical Trial. American Journal of Psychiatry 176, 923–930. 10.1176/appi.ajp.2019.18101123

Denny, R.R., Connelly, K.L., Ghilotti, M.G., Meissler, J.J., Yu, D., Eisenstein, T.K., Unterwald, E.M., 2021. Artificial Intelligence Identified Resilient and Vulnerable Female Rats After Traumatic Stress and Ethanol Exposure: Investigation of Neuropeptide Y Pathway Regulation. Front Neurosci 15. 10.3389/fnins.2021.772946

Duek, O., Korem, N., Li, Y., Kelmendi, B., Amen, S., Gordon, C., Milne, M., Krystal, J.H., Levy, I., Harpaz-Rotem, I., 2023. Long term structural and functional neural changes following a single infusion of Ketamine in PTSD. Neuropsychopharmacology 48, 1648–1658. 10.1038/s41386-023-01606-3

Fleckenstein, A.E., Volz, T.J., Riddle, E.L., Gibb, J.W., Hanson, G.R., 2007. New Insights into the Mechanism of Action of Amphetamines. Annu Rev Pharmacol Toxicol 47, 681–698. 10.1146/annurev.pharmtox.47.120505.105140

Flores, R.J., Cruz, B., Uribe, K.P., Correa, V.L., Arreguin, M.C., Carcoba, L.M., Mendez, I.A., O’Dell, L.E., 2020. Estradiol promotes and progesterone reduces anxiety-like behavior produced by nicotine withdrawal in female rats. Psychoneuroendocrinology 119, 104694. 10.1016/j.psyneuen.2020.104694

Funke, J.R., Hwang, E.-K., Wunsch, A.M., Baker, R., Engeln, K.A., Murray, C.H., Milovanovic, M., Caccamise, A.J., Wolf, M.E., 2023. Persistent Neuroadaptations in the Nucleus Accumbens Core Accompany Incubation of Methamphetamine Craving in Male and Female Rats. eNeuro 10, ENEURO.0480-22.2023. 10.1523/ENEURO.0480-22.2023

Giacometti, L., Barker, J., 2020. Sex differences in the glutamate system: Implications for addiction. Neurosci Biobehav Rev 113, 157–168. 10.1016/j.neubiorev.2020.03.010

Giacometti, L.L., Buck, L.A., Barker, J.M., 2022. Estrous cycle and hormone regulation of stress-induced reinstatement of reward seeking in female mice. Addiction Neuroscience 4, 100035. 10.1016/j.addicn.2022.100035

Hámor, P.U., Knackstedt, L.A., Schwendt, M., 2023. The role of metabotropic glutamate receptors in neurobehavioral effects associated with methamphetamine use. International Review of Neurobiology, volume 168, pp. 177–219. 10.1016/bs.irn.2022.10.005

Kalivas, P.W., LaLumiere, R.T., Knackstedt, L., Shen, H., 2009. Glutamate transmission in addiction. Neuropharmacology 56, 169–173. 10.1016/j.neuropharm.2008.07.011

Kalivas, P.W., Volkow, N.D., 2005. The Neural Basis of Addiction: A Pathology of Motivation and Choice. American Journal of Psychiatry 162, 1403–1413. 10.1176/appi.ajp.162.8.1403

Key Substance Use and Mental Health Indicators in the United States: Results from the 2021 National Survey on Drug Use and Health, 2021.

Kogan, F.J., Nichols, W.K., Gibb, J.W., 1976. Influence of methamphetamine on nigral and striatal tyrosine hydroxylase activity and on striatal dopamine levels. Eur J Pharmacol 36, 363–371. 10.1016/0014-2999(76)90090-X

Krupitsky, E.M., Burakov, A.M., Dunaevsky, I. V, Romanova, T.N., Slavina, T.Y., Grinenko, A.Y., 2007. Single versus repeated sessions of ketamine-assisted psychotherapy for people with heroin dependence. J Psychoactive Drugs 39, 13–9. 10.1080/02791072.2007.10399860

Lewandowski, S.I., Hodebourg, R., Wood, S.K., Carter, J.S., Nelson, K.H., Kalivas, P.W., Reichel, C.M., 2023. Matrix metalloproteinase activity during methamphetamine cued relapse. Addiction Biology 28. 10.1111/adb.13279

Li, Y., Qu, L., Li, N., Wang, Xin, Wang, P., Ge, S., Wang, Xue-lian, 2022. The optimized jugular vein catheterization reinforced cocaine self-administration addictive model for adult male Sprague– Dawley rats. Sci Rep 12, 11711. 10.1038/s41598-022-15833-z

Machado-Vieira, R., Salvadore, G., DiazGranados, N., Zarate, C.A., 2009. Ketamine and the next generation of antidepressants with a rapid onset of action. Pharmacol Ther 123, 143–150. 10.1016/j.pharmthera.2009.02.010

Mark, K.A., Soghomonian, J.-J., Yamamoto, B.K., 2004. High-Dose Methamphetamine Acutely Activates the Striatonigral Pathway to Increase Striatal Glutamate and Mediate Long-Term Dopamine Toxicity. The Journal of Neuroscience 24, 11449–11456. 10.1523/JNEUROSCI.3597-04.2004

Mattson, M.P., 2008. Glutamate and Neurotrophic Factors in Neuronal Plasticity and Disease. Ann N Y Acad Sci 1144, 97–112. 10.1196/annals.1418.005

McGregor, C., Srisurapanont, M., Jittiwutikarn, J., Laobhripatr, S., Wongtan, T., White, J.M., 2005. The nature, time course and severity of methamphetamine withdrawal. Addiction 100, 1320–1329. 10.1111/j.1360-0443.2005.01160.x

McKetin, R., Dawe, S., Burns, R.A., Hides, L., Kavanagh, D.J., Teesson, M., McD. Young, R., Voce, A., Saunders, J.B., 2016. The profile of psychiatric symptoms exacerbated by methamphetamine use. Drug Alcohol Depend 161, 104–109. 10.1016/j.drugalcdep.2016.01.018

Moghaddam, B., Adams, B., Verma, A., Daly, D., 1997. Activation of Glutamatergic Neurotransmission by Ketamine: A Novel Step in the Pathway from NMDA Receptor Blockade to Dopaminergic and Cognitive Disruptions Associated with the Prefrontal Cortex. The Journal of Neuroscience 17, 2921– 2927. 10.1523/JNEUROSCI.17-08-02921.1997

Moriguchi, S., Watanabe, S., Kita, H., Nakanishi, H., 2002. Enhancement of N-methyl-D-aspartate receptor-mediated excitatory postsynaptic potentials in the neostriatum after methamphetamine sensitization. An in vitro slice study. Exp Brain Res 144, 238–46. 10.1007/s00221-002-1039-3

Nagy, E.K., Overby, P.F., Leyrer-Jackson, J.M., Carfagno, V.F., Acuña, A.M., Olive, M.F., 2024. Methamphetamine and the Synthetic Cathinone 3,4-Methylenedioxypyrovalerone (MDPV) Produce Persistent Effects on Prefrontal and Striatal Microglial Morphology and Neuroimmune Signaling Following Repeated Binge-like Intake in Male and Female Rats. Brain Sci 14, 435. 10.3390/brainsci14050435

NIDA. 2021, April 13. What treatments are effective for people who misuse methamphetamine? [WWW Document], n.d.. Retrieved from https://nida.nih.gov/publications/research-reports/methamphetamine/what-treatments-are-effective-people-who-misuse-methamphetamine on 2024, April 27.

Nordahl, T.E., Salo, R., Leamon, M., 2003. Neuropsychological Effects of Chronic Methamphetamine Use on Neurotransmitters and Cognition: A Review. J Neuropsychiatry Clin Neurosci 15, 317–325. 10.1176/jnp.15.3.317

Parsegian, A., See, R.E., 2014. Dysregulation of Dopamine and Glutamate Release in the Prefrontal Cortex and Nucleus Accumbens Following Methamphetamine Self-Administration and During Reinstatement in Rats. Neuropsychopharmacology 39, 811–822. 10.1038/npp.2013.231

Paulus, M.P., Stewart, J.L., 2020. Neurobiology, Clinical Presentation, and Treatment of Methamphetamine Use Disorder. JAMA Psychiatry 77, 959. 10.1001/jamapsychiatry.2020.0246

Pena-Bravo, J.I., Penrod, R., Reichel, C.M., Lavin, A., 2019. Methamphetamine Self-Administration Elicits Sex-Related Changes in Postsynaptic Glutamate Transmission in the Prefrontal Cortex. eNeuro 6, ENEURO.0401-18.2018. 10.1523/ENEURO.0401-18.2018

Perrine, S.A., Sheikh, I.S., Nwaneshiudu, C.A., Schroeder, J.A., Unterwald, E.M., 2008. Withdrawal from chronic administration of cocaine decreases delta opioid receptor signaling and increases anxiety-and depression-like behaviors in the rat. Neuropharmacology 54, 355–364. 10.1016/j.neuropharm.2007.10.007

Reichel, C.M., Chan, C.H., Ghee, S.M., See, R.E., 2012. Sex differences in escalation of methamphetamine self-administration: cognitive and motivational consequences in rats. Psychopharmacology (Berl) 223, 371–380. 10.1007/s00213-012-2727-8

Roth, M.E., Carroll, M.E., 2004. Sex differences in the acquisition of IV methamphetamine self-administration and subsequent maintenance under a progressive ratio schedule in rats. Psychopharmacology (Berl) 172, 443–449. 10.1007/s00213-003-1670-0

Ru, Q., Xiong, Q., Zhou, M., Chen, L., Tian, X., Xiao, H., Li, C., Li, Y., 2019. Withdrawal from chronic treatment with methamphetamine induces anxiety and depression-like behavior in mice. Psychiatry Res 271, 476–483. 10.1016/j.psychres.2018.11.072

Ruda-Kucerova, J., Amchova, P., Babinska, Z., Dusek, L., Micale, V., Sulcova, A., 2015. Sex Differences in the Reinstatement of Methamphetamine Seeking after Forced Abstinence in Sprague-Dawley Rats. Front Psychiatry 6. 10.3389/fpsyt.2015.00091

Saland, S.K., Kabbaj, M., 2018. Sex Differences in the Pharmacokinetics of Low-dose Ketamine in Plasma and Brain of Male and Female Rats. J Pharmacol Exp Ther 367, 393–404. 10.1124/jpet.118.251652

Sarkar, A., Kabbaj, M., 2016. Sex Differences in Effects of Ketamine on Behavior, Spine Density, and Synaptic Proteins in Socially Isolated Rats. Biol Psychiatry 80, 448–456. 10.1016/j.biopsych.2015.12.025

Shrestha, P., Katila, N., Lee, S., Seo, J.H., Jeong, J.-H., Yook, S., 2022. Methamphetamine induced neurotoxic diseases, molecular mechanism, and current treatment strategies. Biomedicine & Pharmacotherapy 154, 113591. 10.1016/j.biopha.2022.113591

Smith, K.J., Butler, T.R., Self, R.L., Braden, B.B., Prendergast, M.A., 2008. Potentiation of N-methyl-d-aspartate receptor-mediated neuronal injury during methamphetamine withdrawal in vitro requires co-activation of IP3 receptors. Brain Res 1187, 67–73. 10.1016/j.brainres.2007.10.019

Su, H., Zhang, J., Ren, W., Xie, Y., Tao, J., Zhang, X., He, J., 2017. Anxiety level and correlates in methamphetamine-dependent patients during acute withdrawal. Medicine 96, e6434. 10.1097/MD.0000000000006434

Suzuki, A., Hara, H., Kimura, H., 2023. Role of the AMPA receptor in antidepressant effects of ketamine and potential of AMPA receptor potentiators as a novel antidepressant. Neuropharmacology 222, 109308. 10.1016/j.neuropharm.2022.109308

Szumlinski, K.K., Lominac, K.D., Campbell, R.R., Cohen, M., Fultz, E.K., Brown, C.N., Miller, B.W., Quadir, S.G., Martin, D., Thompson, A.B., von Jonquieres, G., Klugmann, M., Phillips, T.J., Kippin, T.E., 2017. Methamphetamine Addiction Vulnerability: The Glutamate, the Bad, and the Ugly. Biol Psychiatry 81, 959–970. 10.1016/j.biopsych.2016.10.005

Takashima, Y., Tseng, J., Fannon, M.J., Purohit, D.C., Quach, L.W., Terranova, M.J., Kharidia, K.M., Oliver, R.J., Mandyam, C.D., 2018a. Sex Differences in Context-Driven Reinstatement of Methamphetamine Seeking is Associated with Distinct Neuroadaptations in the Dentate Gyrus. Brain Sci 8, 208. 10.3390/brainsci8120208

Tizabi, Y., Bhatti, B.H., Manaye, K.F., Das, J.R., Akinfiresoye, L., 2012. Antidepressant-like effects of low ketamine dose is associated with increased hippocampal AMPA/NMDA receptor density ratio in female Wistar–Kyoto rats. Neuroscience 213, 72–80. 10.1016/j.neuroscience.2012.03.052

Vuong, S.M., Oliver, H.A., Scholl, J.L., Oliver, K.M., Forster, G.L., 2010. Increased anxiety-like behavior of rats during amphetamine withdrawal is reversed by CRF2 receptor antagonism. Behavioural Brain Research 208, 278–281. 10.1016/j.bbr.2009.11.036

Zanos, P., Gould, T.D., 2018. Mechanisms of ketamine action as an antidepressant. Mol Psychiatry 23, 801–811. 10.1038/mp.2017.255

Zarate, C.A., Manji, H.K., 2008. The Role of AMPA receptor modulation in the treatment of neuropsychiatric diseases. Exp Neurol 211, 7–10. 10.1016/j.expneurol.2008.01.011

Zaytseva, A., Bouckova, E., Wiles, M.J., Wustrau, M.H., Schmidt, I.G., Mendez-Vazquez, H., Khatri, L., Kim, S., 2023. Ketamine’s rapid antidepressant effects are mediated by Ca2+-permeable AMPA receptors. Elife 12. 10.7554/eLife.86022

Zorick, T., Nestor, L., Miotto, K., Sugar, C., Hellemann, G., Scanlon, G., Rawson, R., London, E.D., 2010. Withdrawal symptoms in abstinent methamphetamine-dependent subjects. Addiction 105, 1809– 1818. 10.1111/j.1360-0443.2010.03066.x

